# The genetic and linguistic structures of Abyssinians and their neighbor reveal the historical demographic dynamics and environmental adaptation in the African Horn region

**DOI:** 10.1101/2020.10.21.348599

**Authors:** Demissew Sertse, Tesfaye B. Mersha, Jemaneh Z. Habtewold

**Affiliations:** University of Ottawa, Department of Biology, Ottawa, ON, Canada; University of Cincinnati, Cincinnati Children’s Hospital Medical Center, Department of Pediatrics, Cincinnati, OH, USA; Ottawa Research and Development Centre, Agriculture and Agri-Food Canada, Ottawa, Canada

**Keywords:** Horn of Africa, Ethiopia, ancestry, genetic structure, linguistic variation, haplotype, adaptive loci, geneflow

## Abstract

The African Horn region that includes the Abyssinian is one of the areas in the world that harbor high human genetic diversity manifesting past intermingling of people of different origins attributed to its geographic immediacy to the middle east and being historical trade and religio-cultural hub. Here, we performed a genetic structure analysis of linguistically differentiated populations of Ethiopia, South Sudan, and Somali. To get insight into the genetic landscape of the horn of Africa against the rest of the world, we leverage HapMap SNPs data from Utah residents with Northern and Western European ancestry (CEU), Maasai (MKK), and Yoruba (YRI) and analyzed for genetic admixture and diversity. The genetic and linguistic affiliations mismatch for most Cushitic and Semitic language speakers. The gradients of genetic variations among the different sub-populations within the region show gene-flow directions and past mass population movements. Ethiopians that predominately inhabited the central and northern Ethiopia harbored ~10-15% of CEU admixture. The African horn ancestral line contributed a total of ~20%, 5%, and 2% to MKK, YRI, and CEU, respectively. MKK showed a high genetic diversity comparable to the Ethiopian Cushitic, Semitic, and North Omotic language speakers. Allelic distribution frequencies among the populations at some outlier loci may also provide insight into the adaptations to critical environmental factors such as Malaria.

## Introduction

The Abyssinians in the African Horn archived the ever known length of human history based on hominin fossil recordings so far unearthed (Johanson and White 1979; White et al. 2009). The region is also known for harboring the wider range of the modern human race genetic lineages attributed to gene flow between this region and the rest of the world (Pagani et al. 2012). The ancient kingdoms in the region had expanded their empires that span deep south in Africa and southern Arabia, which perhaps contributed to the demographic flows within their empire and hence genetic admixture(Breton et al. 2014). The old Christian scripts circuitously described the diversity of the Abyssinian inhabitants; while Jeremiah 13:23 mentions the unique color of Ethiopians, Ezkiel30:5 lists Ethiopians among the mingled races. Abyssinia was a trade hub (Krzemnicki et al. 2019; Sernicola and Phillipson 2011) and destination of people of different origins (Phillipson 1993). This has formed the noticeably observed mosaic appearances of the people and religio-cultural diversity. Owing to their geographic proximity, the northern Abyssinian highlanders are immensely influenced by the Middle Eastern, which goes faint to the south and eventually dominated by indigenous African customs resulting in a gradient pattern of variations.

The genetic structure of Ethiopians was previously shown to follow the linguistic variation pattern (Pagani et al. 2012). However, the historical mass migrations and assimilation events of people may lead to incongruent genetic and linguistic stratification. Within the current boundary of Ethiopia, there were important migrations of people. Among others, the earlier southward Judeo-Christian northern highlanders’ expansion (Nelson and Kaplan 1981) which perhaps resulted in spread of people as far south as the southern extreme of the continent Africa (Breton et al. 2014) and the later south-north movement as the case of the 16^th^ century Oromo mass movement can be mentioned (Nelson and Kaplan 1981). While it is expected that these kinds of movements can result in significant genetic admixture, the linguistic and cultural assimilation of certain people with their intact original genetic makeup cannot be undermined. i.e, some community in the south that currently share culture and language with the north highlanders may have different ancestral roots, and some with disparate linguistic families can genetically be closely related. This applies to the community currently living in the northern highlands. The extant Zay community that has been confined in the Lake Ziway islands with their survived original culture and language (Jordan et al. 2011) in the Oromo dominant Ziway area can be one of the indications.

On the other hand, many cultures and languages that had existed before the Oromo movement have eventually been assimilated and led to extinctions. At the same time, those original settlers may still live in their intact genetic makeup. This is true for many communities currently speaking Amharic in the north. Perhaps the current dominant Amharic speakers are the Agew and other early inhabitants of the region (Bender 1971). As such the term Amhara was ascribed to Abyssinian highlanders regardless of their background (Clark 1950) which might reflect the argument of some scholars who claim the Origin of the term from the Heberew ‘am hare’; ((am = people, Hare = mountain). In line with decedents of different origins living together, high genetic diversity and admixture are expected in Ethiopians lineage (Pagani et al. 2012).

The current molecular techniques are opportunities to disentangle the existing linguistic as well as cultural mingling and the genetic structure of a society in a given geography. The Ethiopian genetic structure has been a subject for a couple of studies (Llorente et al. 2015; López et al. 2019; Pagani et al. 2012; van Dorp et al. 2015). Because of Ethiopia’s geographical position between Africa and Eurasia (Europe and Asia) and expansion point for modern human “out of Africa” or “back to Africa”, it has a unique population structure that has attracted scientists (Lawson et al. 2018; Llorente et al. 2015). In this study, we applied genetic fingerprints using SNPs information to unravel the historical evidences on population admixture, geneflows, and linguistic family in Ethiopia and across the horn of Africa.

## Materials and methods

We downloaded the Ethiopian genetic data from http://mega.bioanth.cam.ac.uk/data/Ethiopia/ stored by (Pagani et al. 2012). This data included 235 individuals genotyped for ~one million SNP each. The individuals were from a total of 12 ethnic groups, of which two: South Sudanese and Somali from mainland Somalia were outside the Ethiopian border. The data were filtered to SNPs of no missing point with a 10% minor allele frequency (MAF). To perform a genome scan and compare result with no missing dataset (144k), another dataset of SNPs of 95% call rate, with 10% MAF was prepared by pruning out one of the pair SNP accounting linkage disequilibrium (LD) of r^2^>0.8 using Plink 2.0 (Purcell et al. 2007).

To provide insights into the genetic landscapes of Ethiopians against others, SNP data of three populations with a total of 496 individuals : Utah residents with Northern and Western European ancestry (CEU, n=162), Maasai (MKK, n=171), and Yoruba (YRI, n=163) were downloaded from Hapmap3 site ftp://ftp.ncbi.nlm.nih.gov/hapmap/genotypes/hapmap3_r3/hapmap_format/consensus/ and merged with original Ethiopian genetic dataset. Only SNPs with 95% and above call rate and common in all populations were filtered from the merged data for further analyses.

### Genetic structure, admixture, and variation analyses

To estimate the appropriate number of subpopulations (K) for the Ethiopian genetic dataset, principal component analysis (PCA) was performed, based on the maximum 30 principal components (PCs), using the R package pcadapt version 4 (Privé et al. 2020). The appropriate number of ancestral subgroups (K) was defined following the PC before the last curve of the on screeplot, as described by (Luu et al. 2017). The estimate of appropriate K for the merged dataset was performed using a cross-validation technique (Alexander and Lange 2011) implemented in the program ADMIXTURE (Alexander et al. 2009).

The neighbor-joining (NJ) phylogenetic analysis was performed using TASSEL v 5 (Bradbury et al. 2007) and visualized using a web-based software Interactive Tree of Life (ITOL) (Letunic and Bork 2016). Since the estimated appropriate K for the Ethiopian dataset appeared difficult to define among three marks: K=5, 8, and 12, to show the entire span, each individual’s ancestral coefficients were calculated from K2 to K12 using sNMF (Frichot et al. 2014) with snmf function implemented in R package LEA (Frichot and François 2015). For the merged dataset, the ancestral coefficients of individuals were computed while in the meantime generating the cross-validation error (Alexander and Lange 2011) from K2-K20 using ADMIXTURE (Alexander et al. 2009). However, to show the admixture, only Ks including estimated appropriate K (Ke), Ke±1 with the lowest cross-validation error, K2, K16 and K20 were presented. The results of ancestral estimates from both Ethiopian and merged datasets were summarized using Clustering Markov Packager Across K (CLUMPAK) http://clumpak.tau.ac.il/distruct.html. Principal component analyses were performed for the two datasets separately at their corresponding best K and PCs 1-3 from PCA of each dataset were plotted to show the clustering patterns. To determine the genetic and linguistic structure of the Ethiopian populations, phylolingustic tree data of Ethiopian and other languages in the African horn region were extracted from world phylolingustic tree downloaded from https://osf.io/cufv7/files/.

Pairwise genetic differentiation among the Ethiopian groups was computed based on the 144k dataset using an R package diveRsity (Keenan et al. 2013) following F_ST_ algorism by (Weir and Cockerham 1984). F_ST_ of each locus was also computed, and the haplotypes were generated based on 100 top high F_ST_ SNPs using web tool Sniplay (Dereeper et al. 2011). The topmost ten high-frequency haplotypes were selected and mapped on the geographic coordinates that fairly represent the habitation of the populations. Geographic coordinates partly obtained from Pagani et al. (2012), and for the rest, coordinates of locations best represent each population was retrieved from Google map. The genetic differentiation and diversity analyses for the merged dataset was performed using Genepop in R(Rousset 2008) following the procedures by Weir and Cockerham (1984). The gene diversity analysis was performed for each individual and population.

### Genome scan and potential adaptive loci

To identify loci underlying local adaptations from Ethiopian dataset, genome scan was performed based on both 144k and 95% call pruned datasets using an R package pcadapt v.4.0 (Luu et al. 2017) at estimated ancestral populations K=5, K=8 and K=12. The numbers of appropriate ancestral populations were inferred from screeplot generated by the pcadapt. F_ST_ based outlier loci analysis was also performed for the Ethiopian 144k dataset using the R package LEA (Frichot and François 2015) for Ks mentioned above (K=5,8,12). Similar analysis was performed based on the merged dataset using pcadapt in R (Luu et al. 2017). Top 20 outlier SNPs were recovered and further explored in databases for their clinical or other associated relevancies. Allele frequencies of five most consistent loci from the Ethiopian dataset were calculated for each population. To get insight into worldwide allelic distribution of loci associated with a known environmental factor or clinical relevance, the allele frequencies of these loci were also calculated in the three additional populations: CEU, MKK, and YRI.

## Results

### *Genetic structure* and admixture

All the genetic structure analyses: Admixture, PCA, and NJ show a similar pattern (Fig 1A-E). The Afar, Amhara, Tigray, and Majority of Oromo folded in the same cluster. The NJ indicated that individuals from different ethnic groups were more closely related than individuals of the same ethnic class. Afar took a separate sub-clade under the major clade that embraced Amhara, Tigray, and majority Oromo. A significant number of Oromo individuals were also clustered intermingled with Wolayita. Anuak and south Sudanese indistinctly dropped in same cluster, while Ethiopian Somali and Somali from mainland Somalia were distinguishably sub-grouped. Although Ari blacksmith and Ari cultivators appeared relatively closely related, they consistently held a separate cluster. The admixture analysis showed that most of the Ari blacksmiths were admixed with Ari cultivator. Admixtures of Ari cultivator were also seen in most of the Wolayita and Some Oromo individuals. A higher proportion of the Amhara-Tigrye_Oromo majority’s admixtures were observed in Ethiopian Somali than the Somali from mainland Somalia (Fig 1A). The first two PCs that accounted for over 60% (Fig 1B) total variance, clustered the individuals in five major groups: Amhara, Tigray, Afar, Oromo, Wolayita and the two Somali (G1); Anuak and South Sudan (G2), Gumuz (G3) and two Ari separately (G4 &G5) (Fig 1B-D).

**Figure 1.**
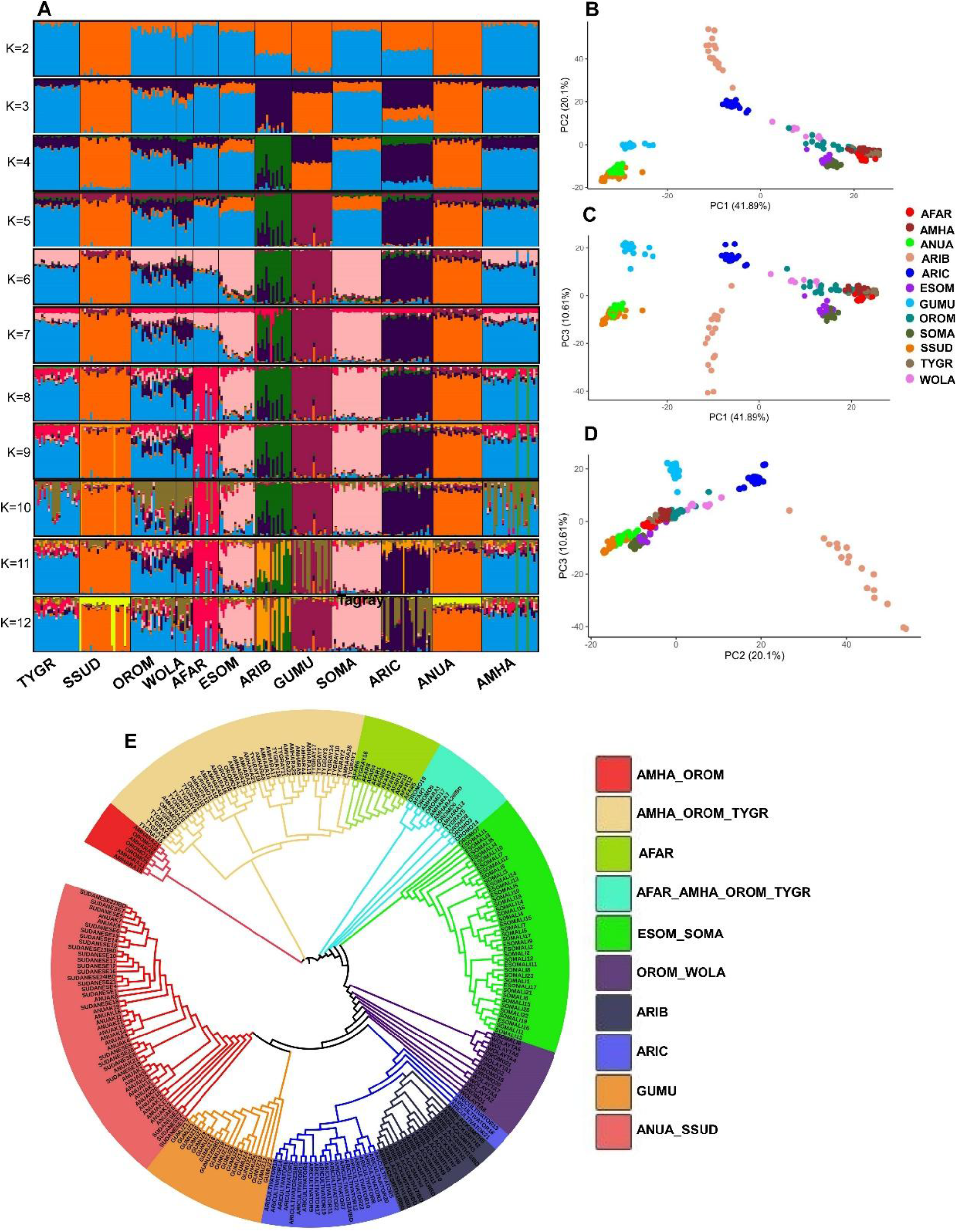
Genetic structure of Ethiopian and people of the horn of Africa. **A**) Genetic admixture from K=2-12; **B-D**) pca of pc1&2, PC1&3, PC2&3 respectively and **E**) NJ Phylogenetic tree.

The highest differentiation was computed between Ari Blacksmith and South Sudanese. Ari Blacksmith appeared deviant in that its smallest differentiation was scored with Ari Cultivator. On the other hand, the smallest differentiation was calculated between Ethiopian Somali and Somali from mainland Somalia (F_ST_=0.0008), followed by between Tigray and Amhara (F_ST_=0.0011). The differentiation among the G1 population was low, where the highest differentiation (F_ST_=0.012) was computed for Somali from mainland Somali and Wolayita (Table 1). The highest and the lowest expected heterozygosity was calculated for Wolayita and South Sudan, respectively. The populations in major group 1 (G1) in general scored higher heterozygosity than the other groups.

**Table 2.**
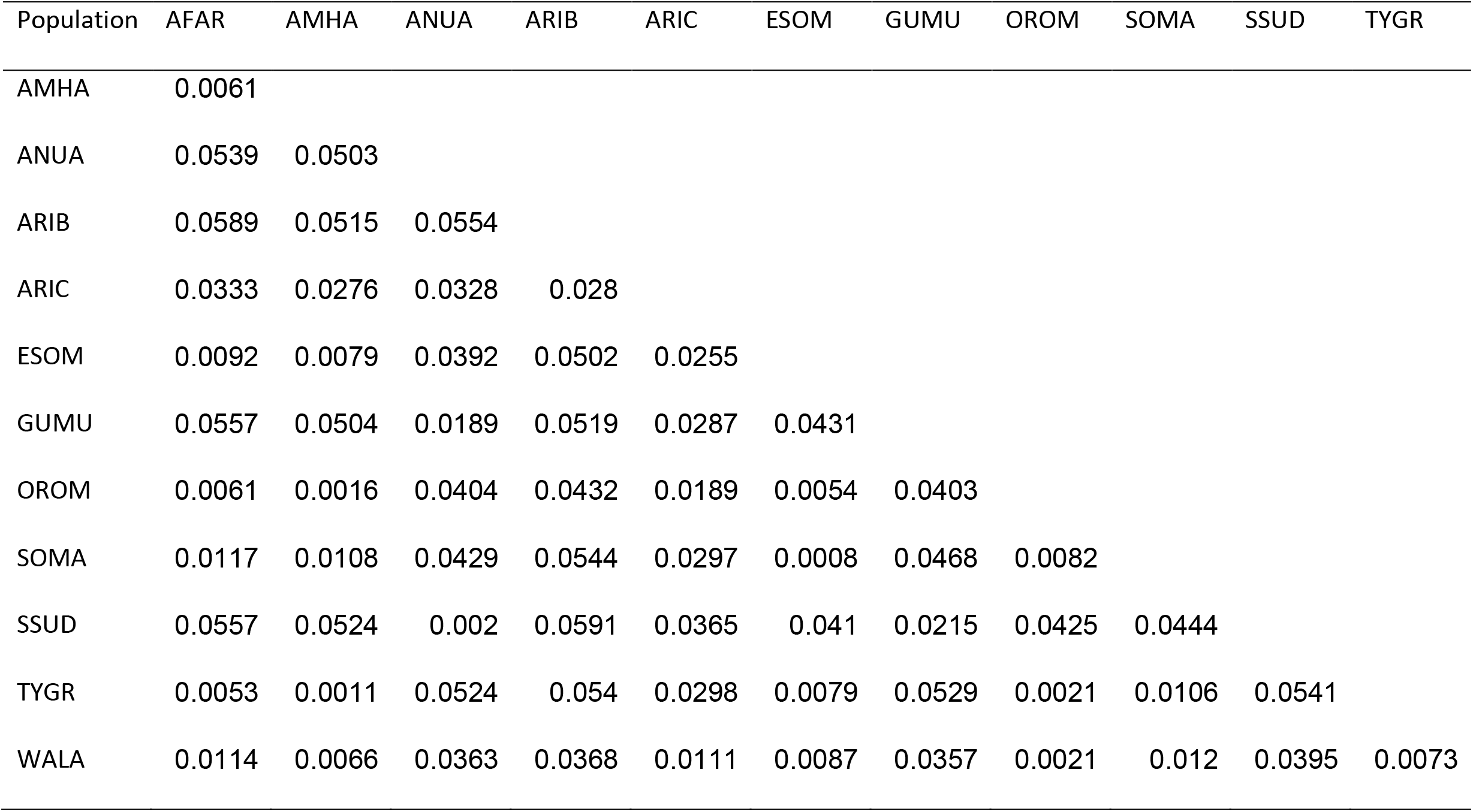
population differentiation among Ethiopian and their Neighbor

| Population | AFAR | AMHA | ANUA | ARIB | ARIC | ESOM | GUMU | OROM | SOMA | SSUD | TYGR |
| --- | --- | --- | --- | --- | --- | --- | --- | --- | --- | --- | --- |
| AMHA | 0.0061 |  |  |  |  |  |  |  |  |  |  |
| ANUA | 0.0539 | 0.0503 |  |  |  |  |  |  |  |  |  |
| ARIB | 0.0589 | 0.0515 | 0.0554 |  |  |  |  |  |  |  |  |
| ARIC | 0.0333 | 0.0276 | 0.0328 | 0.028 |  |  |  |  |  |  |  |
| ESOM | 0.0092 | 0.0079 | 0.0392 | 0.0502 | 0.0255 |  |  |  |  |  |  |
| GUMU | 0.0557 | 0.0504 | 0.0189 | 0.0519 | 0.0287 | 0.0431 |  |  |  |  |  |
| OROM | 0.0061 | 0.0016 | 0.0404 | 0.0432 | 0.0189 | 0.0054 | 0.0403 |  |  |  |  |
| SOMA | 0.0117 | 0.0108 | 0.0429 | 0.0544 | 0.0297 | 0.0008 | 0.0468 | 0.0082 |  |  |  |
| SSUD | 0.0557 | 0.0524 | 0.002 | 0.0591 | 0.0365 | 0.041 | 0.0215 | 0.0425 | 0.0444 |  |  |
| TYGR | 0.0053 | 0.0011 | 0.0524 | 0.054 | 0.0298 | 0.0079 | 0.0529 | 0.0021 | 0.0106 | 0.0541 |  |
| WALA | 0.0114 | 0.0066 | 0.0363 | 0.0368 | 0.0111 | 0.0087 | 0.0357 | 0.0021 | 0.012 | 0.0395 | 0.0073 |

### High F_ST_ SNPs and haplotype distribution

The pattern of distribution of the top 100 high F_ST_ SNPs leaned to go with the major groups of the populations (Fig 2). South Sudanese, Anuak and Gumuz were dominated by homozygous top 100 high F_ST_ SNPs. Whereas, populations under major group 1 had a higher frequency of heterozygous loci while the two Ari harbored approximately equal frequency of the homozygous and heterozygous (Fig 2A).

**Figure 2.**
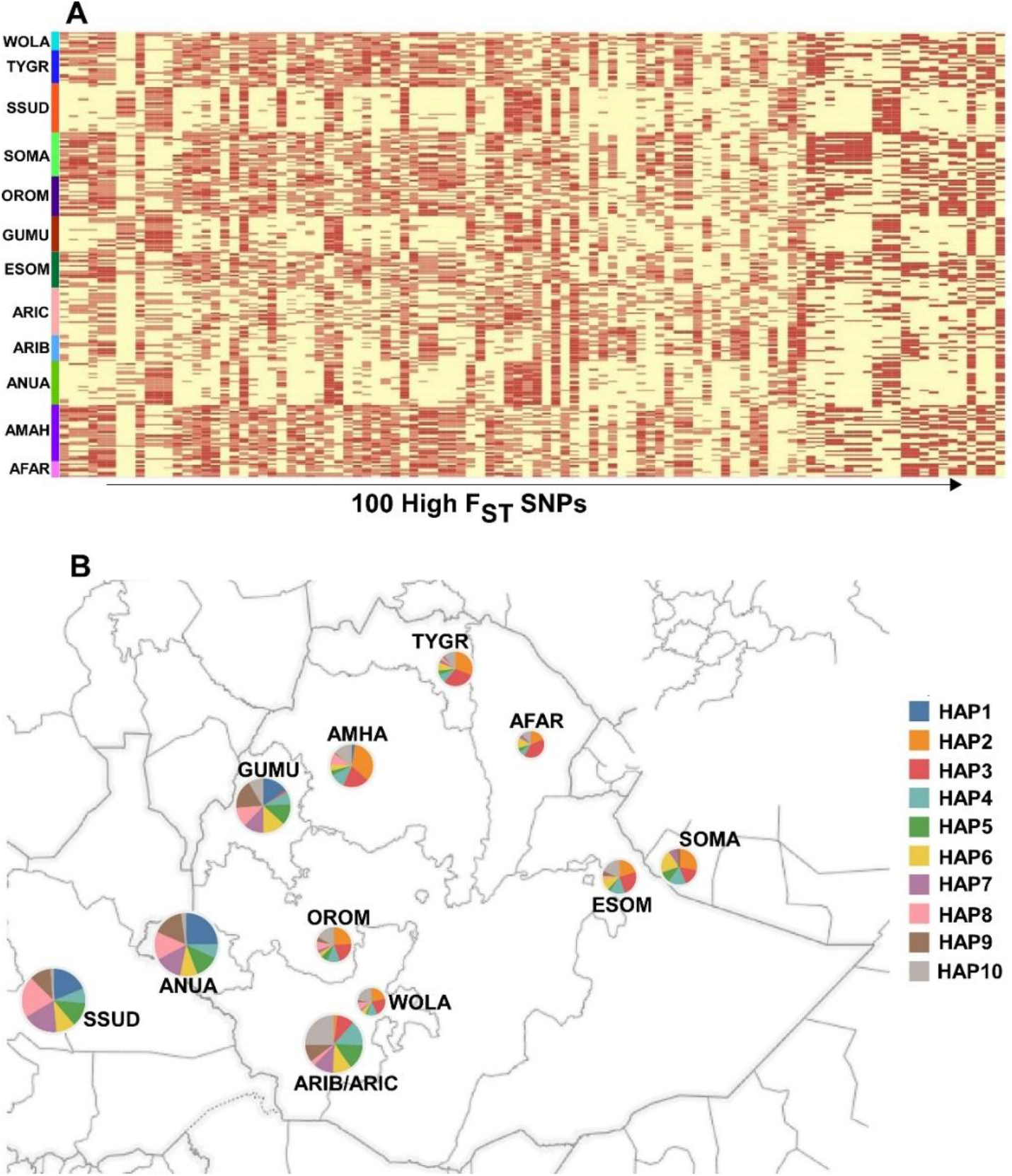
Distribution of top 100 high F_ST_ SNPs and top 10 high frequency haplotypes from the 100 high F_ST_ SNPs; **A**) Top 100 high F_ST_ SNPs, the light yellow indicates dominantly homozygous while the brown indicates heterozygous dominant; Each color of vertical line on the left indicates the sample range of the population it refers to; **B**) The geographic and population distribution of the top 10 high frequency haplotypes; each color in a pie represent a haplotype; the size of the circles indicate the number of individuals in a population.

Of the top ten frequently observed haplotypes generated from the top 100 high F_ST_ SNPs, two haplotypes HAP2 and HAP3 were absent in major group 2 and 3, while they appeared dominantly in all populations of the other groups, where they occurred in more than 50% of individuals in each AFAR, Tigray and Amhara population (Fig 2B). On the other hand, HAP1 that occurred at substantially high frequency in populations under major groups 2 and 3 was absent in the rest populations except in Amhara where it was observed scantly. HAP10, which was the most dominant haplotype in the two Ari populations, spread in all of the populations except in Somali from the mainland Somalia. The Somali from mainland Somalia also lacked HAP8, a haplotype which is also absent in Afar. Afar and Somali each harbored seven out of the ten haplotypes, where Afar privately lacked HAP9 (Fig 2B).

### Genetic and linguistic structure disparity

The phylolingustic tree showed Semitic languages deviant from the rest of the country’s linguistic families, which was against the close genetic relationship of populations under major group 1 that enclosed the Semitic, Cushitic, and north Omotic (Wolayta) languages speakers (Fig 3A-B, 1A-E). This disparity especially was strong in the Cushitic language speakers Afar and the majority of the Oromos and the Semitic speakers Amhara and Tigray, which all were genetically clustered together and showed low genetic differentiation. The pattern of haplotype distribution also showed mismatch with the linguistic structure, where all the Omotic, Semitic and Cushitic speakers shared unique haplotypes that were absent in Nilotic speakers (South Sudanese, Anuak and Gumuz) despite some Omotic and Cushitic languages were relatively in close geographic distance with Nilotic than Semitic (Fig 3 and 2).

**Figure 3.**
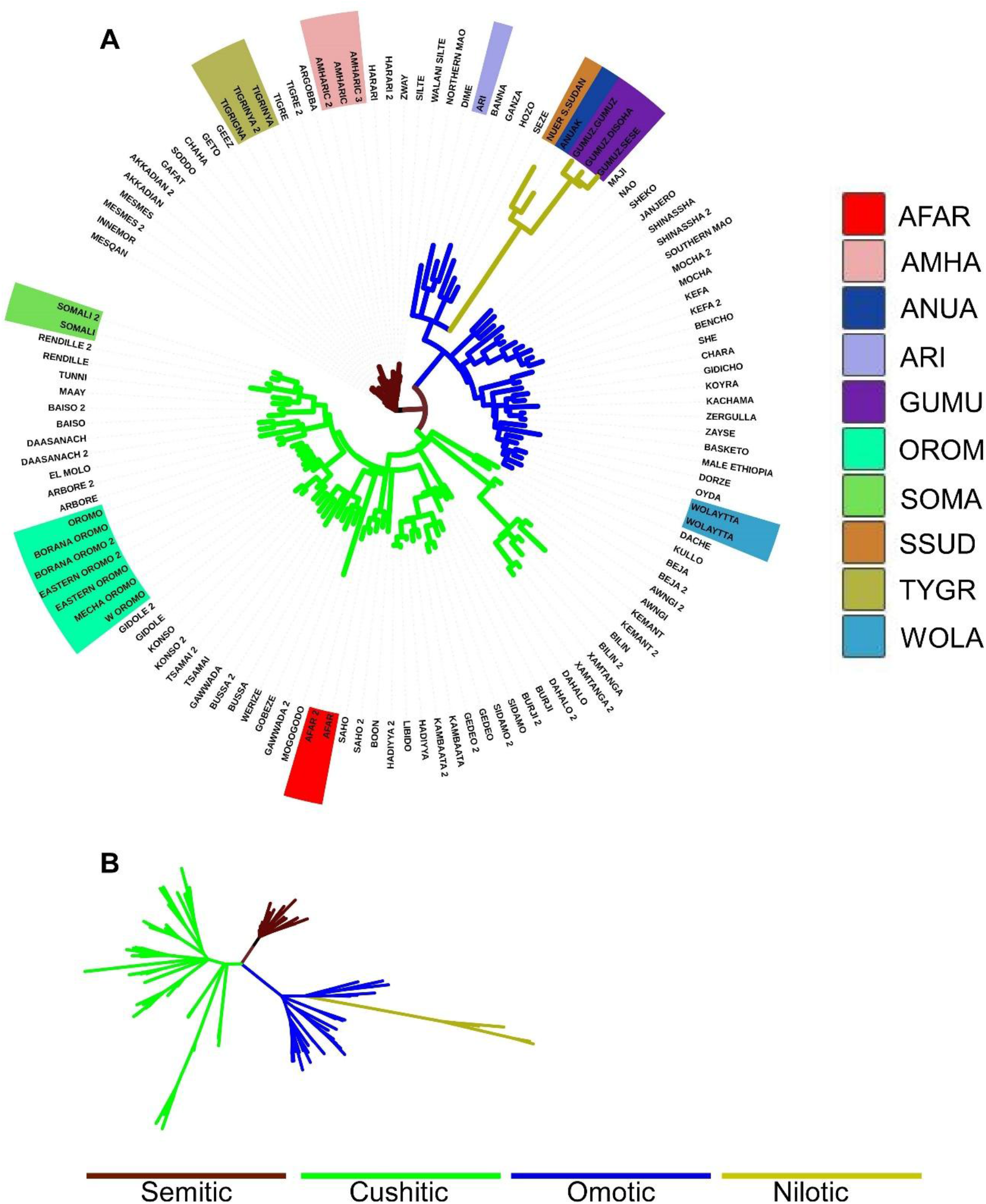
Phylolingustic structure of Ethiopians and the African horn; **A**) phylolingustic relationship of languages in Ethiopia and the vicinity, shaded languages represent the languages of the population targeted in this study; **B**) unrooted topological view of the phylolingustic structure, the colors of the branches in both a and b indicate the major linguistic families.

#### Genetic structure of Ethiopians against others

A total of ~110K SNPs were extracted from the merged data based on the criteria bi-allelic SNP with 95% and above call rate for all merged dataset analyses. The appropriate number of ancestral populations (K) for the merged data that included a total of 15 populations, including the 10 Ethiopian, South Sudanese, and Somali from mainland Somali in Ethiopian dataset and additional three: CEU, MKK and YRI estimated to be seven (Fig 4A). However, K=6 showed a more clear clustering than K=7 (Fig 4B). Afar, Amhara, Oromo, and Tigray under major group 1 showed over 10% admixture from CEU, while those under major group 2 and 3 were noticeably admixed with MKK and YRI. The MKK harbored a higher admixture from CEU than both Ethiopian Somali and the Somali from the mainland Somali (Fig 4B, 5). The unique ancestral line of Ari Blacksmith was distributed in all Ethiopian populations (Fig 4B, 5). The PCA and NJ analyses showed a similar clustering pattern consistent with the Admixture analysis (Fig 4C-F). Generally, MKK is closely related to Ethiopians

**Figure 4.**
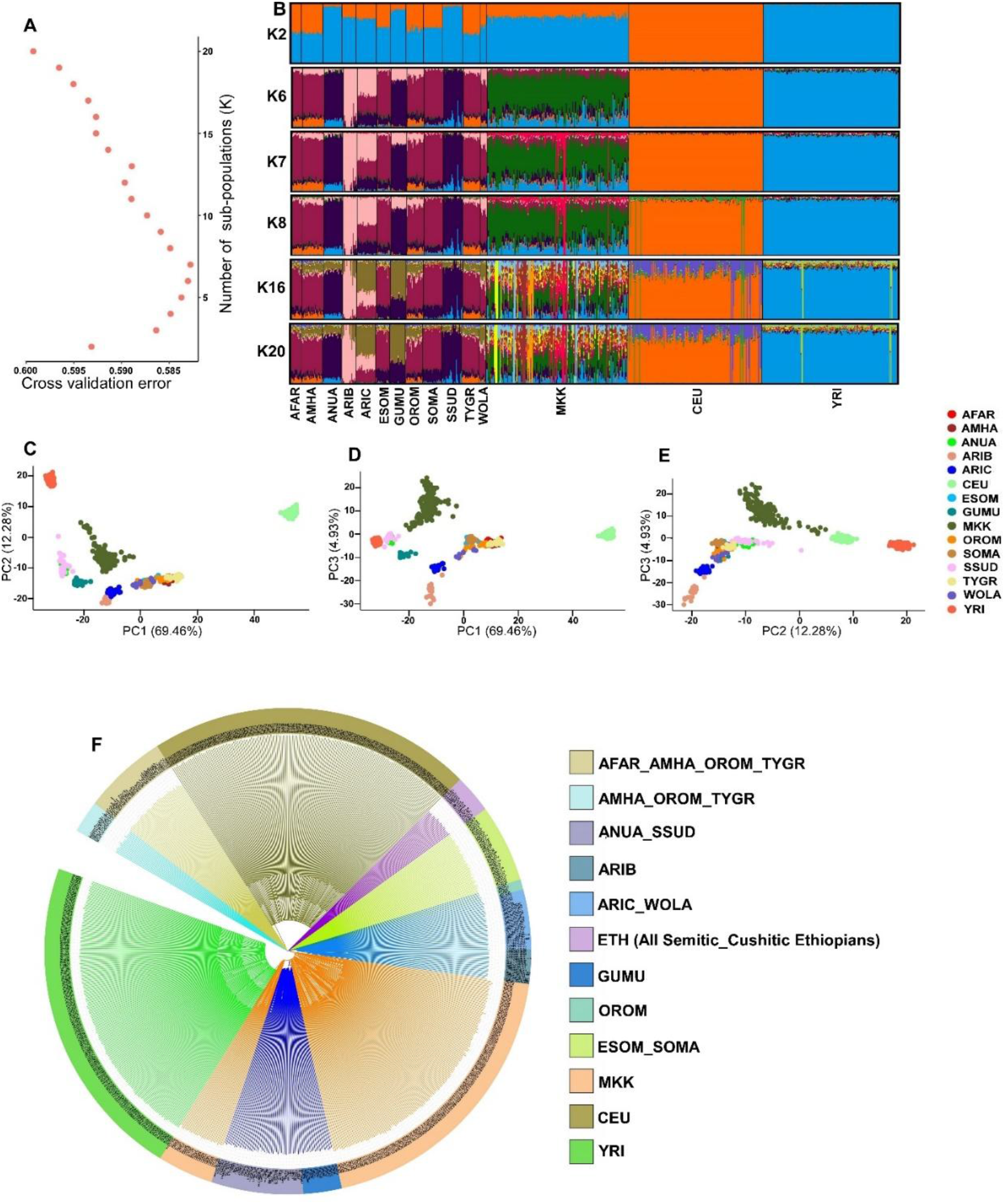
Genetic structure of Ethiopians against others; **A**) graphical presentation of the cross validation errors of K2-20 indicating K=7 the lowest as appropriate number of ancestral populations; **B**) genetic admixture among the populations; **C-E**) clustering of the populations based on PC1-PC3 of the PCA; **F**) NJ phylogenetic trees for the target individuals of all populations.

**Figure 5.**
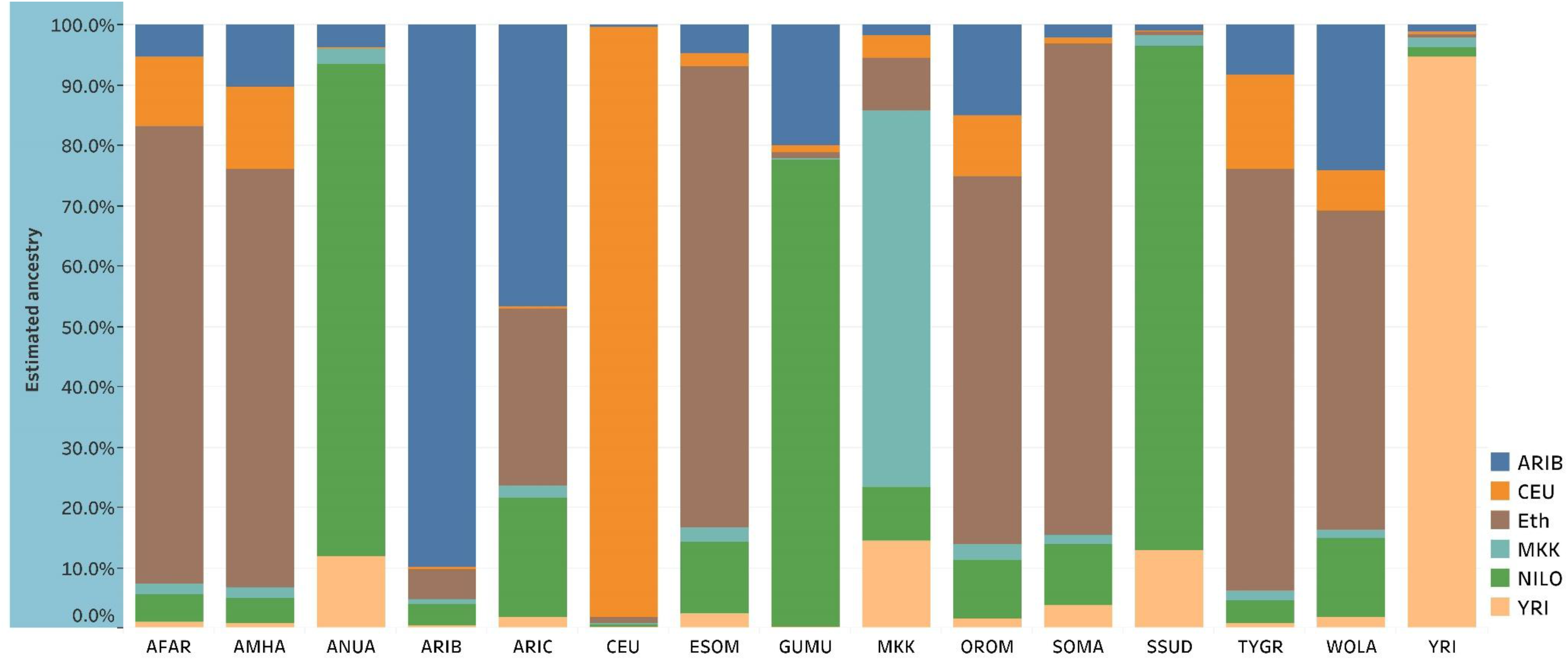
Average estimates of admixture among the ancestral groups: ARIB=Ari Blacksmith; CEU=Europe; Eth=Ethiopians (major group1); MKK=Maasai, NILO= Nilotic; YRI=Yoruba.

Consistent with the genetic structure, a low genetic differentiation was computed between Ethiopians and MKK, the highest and the lowest with Ari Blacksmith (F_ST_=0.04387) and the Oromo (F_ST_ =0.01869), respectively. Likewise, CEU and YRI were remotely related to most Ethiopians, where the lowest and the highest differentiation were observed with Tigray (F_ST_=0.05285) and with Anuak (F_ST_=0.1526), respectively. The highest differentiation of YRI was with Afar (F_ST_=0.07316) (Table 2).

**Table 2.** population differentiation among Ethiopian and other populations.

| Population | AFAR | AMHA | ANUA | ARIB | ARIC | CEU | ESOM | GUMU | MKK | OROM | SOMA | SSUD | TYGR | WOLA |
| --- | --- | --- | --- | --- | --- | --- | --- | --- | --- | --- | --- | --- | --- | --- |
| AMHA | 0.006 |  |  |  |  |  |  |  |  |  |  |  |  |  |
| ANUA | 0.054 | 0.050 |  |  |  |  |  |  |  |  |  |  |  |  |
| ARIB | 0.058 | 0.051 | 0.053 |  |  |  |  |  |  |  |  |  |  |  |
| ARIC | 0.032 | 0.027 | 0.031 | 0.027 |  |  |  |  |  |  |  |  |  |  |
| CEU | 0.066 | 0.061 | 0.153 | 0.138 | 0.114 |  |  |  |  |  |  |  |  |  |
| ESOM | 0.009 | 0.008 | 0.039 | 0.049 | 0.024 | 0.080 |  |  |  |  |  |  |  |  |
| GUMU | 0.056 | 0.050 | 0.018 | 0.050 | 0.028 | 0.149 | 0.042 |  |  |  |  |  |  |  |
| MKK | 0.028 | 0.025 | 0.020 | 0.044 | 0.023 | 0.104 | 0.020 | 0.028 |  |  |  |  |  |  |
| OROM | 0.006 | 0.002 | 0.040 | 0.042 | 0.018 | 0.069 | 0.006 | 0.040 | 0.019 |  |  |  |  |  |
| SOMA | 0.011 | 0.011 | 0.042 | 0.053 | 0.028 | 0.082 | 0.001 | 0.046 | 0.022 | 0.008 |  |  |  |  |
| SSUD | 0.056 | 0.052 | 0.002 | 0.056 | 0.035 | 0.154 | 0.041 | 0.021 | 0.021 | 0.042 | 0.044 |  |  |  |
| TYGR | 0.005 | 0.001 | 0.053 | 0.053 | 0.029 | 0.058 | 0.008 | 0.053 | 0.026 | 0.002 | 0.011 | 0.054 |  |  |
| WOLA | 0.011 | 0.006 | 0.035 | 0.037 | 0.011 | 0.083 | 0.008 | 0.035 | 0.017 | 0.003 | 0.011 | 0.039 | 0.007 |  |
| YRI | 0.073 | 0.070 | 0.031 | 0.072 | 0.053 | 0.158 | 0.060 | 0.046 | 0.028 | 0.060 | 0.062 | 0.031 | 0.072 | 0.055 |

Ethiopians in major group 1 appeared superior in their gene diversity where the highest individual and population gene diversity was computed for the Ethiopians Wolayita (G= 1.3259) and Oromo (G=1.3148), respectively. The Ethiopian major groups 2 & 3, Ari Blacksmith, CEU, and YRI were with approximately similar low genetic diversity. The Ari Blacksmith was with the lowest population gene diversity (G=1.1996). Both individual and population gene diversities for MKK were comparable with the Ethiopians in major group 1(Fig 6).

**Figure 6.**
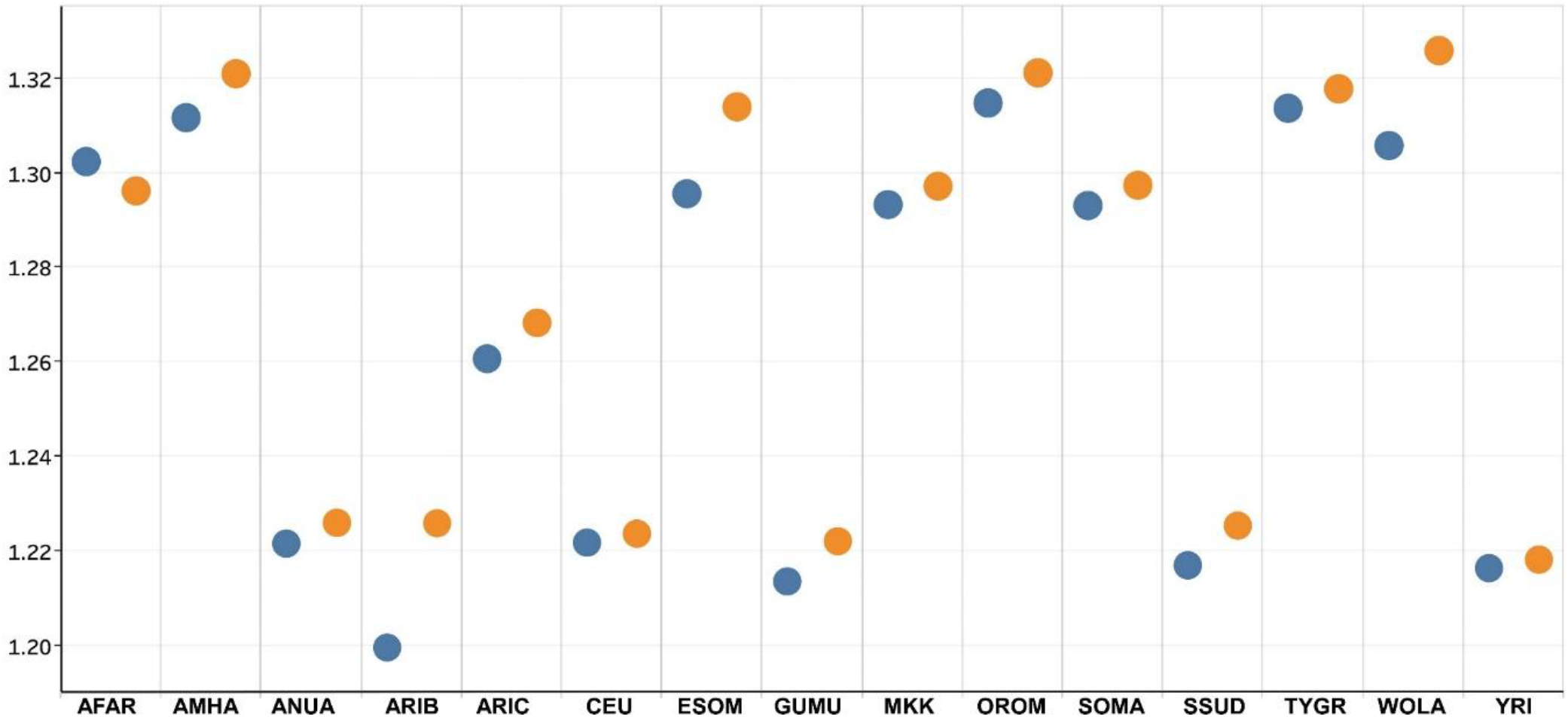
Genetic diversity of Ethiopians against others. The blue and orange circles indicate genetic diversity within population and individual respectively.

### Genome scans for adaptive loci

A total of five outlier loci were detected by a minimum of three times in the two Ethiopian datasets (144k and pruned data) following genome scan at three Ks (K=5, 8, and 12) (Fig S1). Allelic frequencies at these loci largely differed among major groups. One of these loci was a reportedly malaria resistance-associated rs10900588 (Gouveia et al. 2019). The Nilotics: Anuak, Gumuz, and South Sudanese harbored the G allele at this SNP locus exceptionally at high frequencies. This allele was also observed in the South Omotic: the Aries (ARIB & ARIC) in relatively higher frequency than in the remaining populations (Fig 7). The frequency of G allele at this SNP in YRI was comparable with its frequency in Ethiopian Nilotics and the South Sudanese. MKK harbored G at a frequency between the Nilotic and the South Omotic (Fig S4). SNPs associated with some certain cancer and diabetics were also detected in the merged data (Table S1)

**Figure 7.**
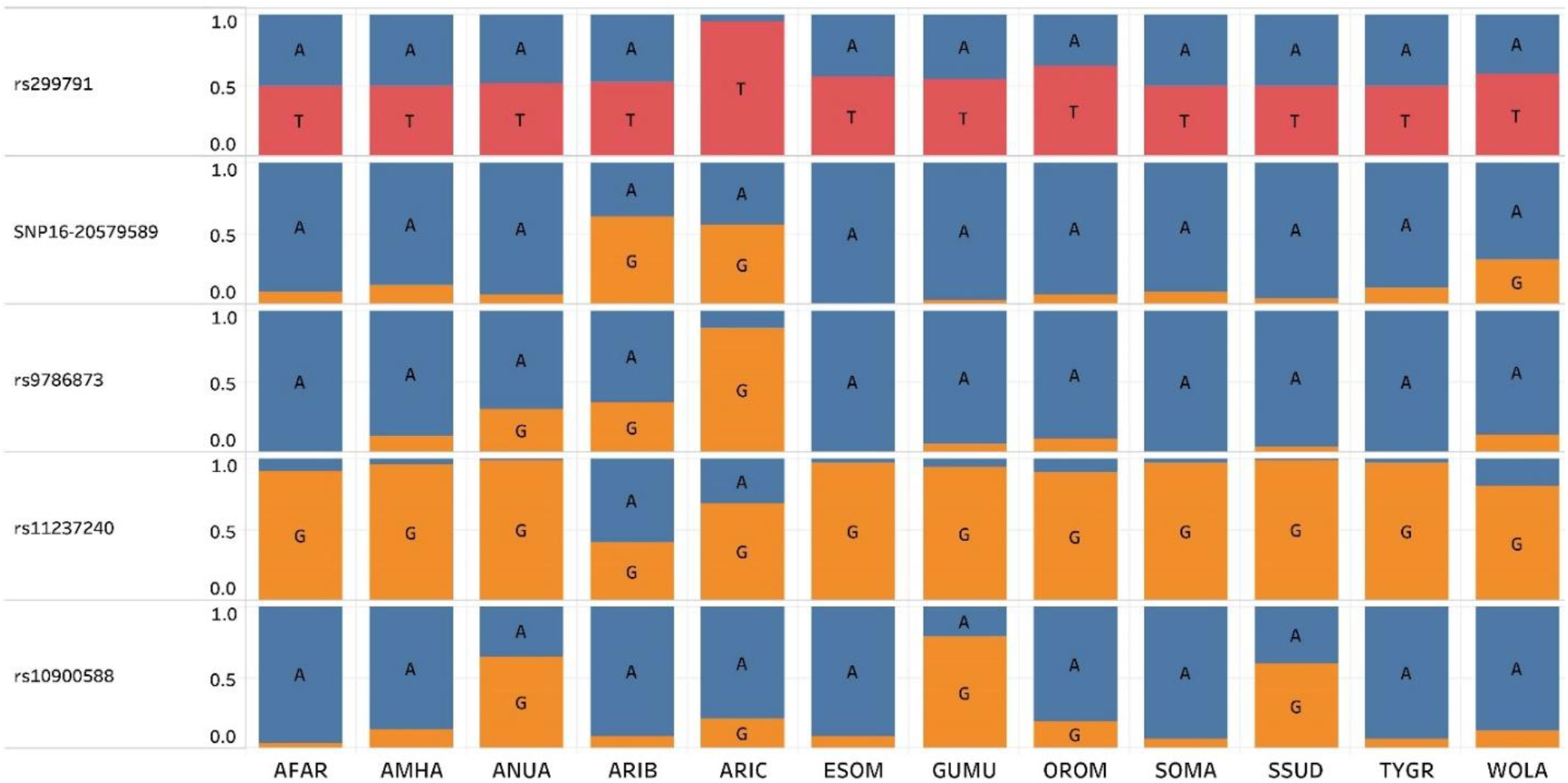
Allelic distribution of to five potential adaptive loci most detected at K 5, 8 and 12 based on different data sets and genome scan methods.

## Discussion

### Ethiopian genetic structure disparity with the current linguistic and ethnic status

Both the genetic and linguistic structures of populations provide insights into the historical contacts of populations. Hence, integrating genetic and linguistic data have become one of the approaches in tracing the human populations’ divergence and admixture histories (Duda and Zrzavý 2019; Kusuma et al. 2016; Li et al. 2020; Thouzeau et al. 2017). However, the linguistic and genetic relationship may not follow a similar pattern since the former is inherited, while the latter is a learned constituent of humans(Duda and Zrzavý 2019; Kusuma et al. 2016). As a result, linguistically related populations may not have a genetic relation and vice versa. This study shows a partial disparity between the genetic and linguistic structure of the Ethiopians. The Ethiopian population genetic structure uncovers the outcomes of historical population movements and assimilation incidences that the linguistic structure has masked.

The results in this study are likely attributed to major population migration and assimilation events; of which the two are the earlier north-south spread since ~5000 years (Harrower et al. 2010; Lesur et al. 2014) and the 16^th^ century south to north Oromo expansion (Hassen 2015). The consistent and undistinguishable meshing up of the Afar, Amhara, Tigray, and majority Oromo in the same cluster against the wide linguistic distance possibly manifests the assimilations of people by others. The close genetic relationship of the Amhara and Tigray evidently acknowledges the widely known strong historical bond between the two communities (Levine 1974).

Unlike the Tigray and Amhara, the strong genetic relationship of most Oromos, especially with the Amharas, is likely associated with the assimilation Amharas and other related highlanders by the Oromo during the post 16^th^ century Oromo migration period. Before the 16^th^ century mass expansion of the Oromos from south to the north, there had reasonably been less contacts between Oromos and the Northerners: Amhara, Tigray, and Afar, at least for some centuries. The Oromo movement was typically characterized by its intense assimilation processes via the Oromo Geda system that changed many communities into Oromo where the Oromos concurred (Hassen 1983). The assimilated groups were recognized from the ‘pure Oromos’ as ‘Gabbaro’ or ‘Gebare’ (Baxter et al. 1996).The ‘Gabbaros’ were renamed after one of the leaders of the ‘pure Oromo’ line abandoning their original names that reflects their biological paternal line (Baxter et al. 1996). Some of these ‘Gabbaros’, particularly the Shewa highlanders, continued to live in their own land in their intact niche (Hassen 1983). The majority of the current Oromo language speakers of the Shewa highlanders are assumed to be decedents of the Shewa Christian community (Taddesse 1968) speaking Amharic, Guraghe, particularly the Soddoo Guraghe (Fig 3), and other languages of mostly Semitic family (Jordan et al. 2011; Ketema 2015; Taddesse 1968). The fingerprints of these communities, such as at Wonchi, Ziqualla, Asebot, and Adadi Mariam have still survived via their old churches and monasteries while the language of the local community of each has eventually been changed in into Oromo language (Fleming 1976; Ketema 2015). The Zay language by the community who are confined in islands of the Zeway Lake is now the only known living pre-Oromo languages in the current Oromo occupied districts of Shewa (Jordan et al. 2011).

The clustering of the Oromos with different population groups and relatively high heterozygosity also is plausibly attributed to the 16^th^-century Oromo expansion and subsequent assimilation and intermingling with others. The clustering of some Oromo individuals with Wolayita can either be post- or pre-Oromo expansion related contacts. The historical origin of Oromo is strongly believed to have been from a region close to Wolayita, where its first expansion record was traced crossing the Galana Sagan River that flows from its source close to Lake-Chamo, a place between Wolayita and Gamo to towards south east, to Lake Chew Bahir (L. Stafanie) (Lewis 1966). This may give us hits that the original Oromos might be closely related to one of the Omotic people. The relatively higher percentage of admixture from Ari-cultivator (Fig 1A) may also strengthen this argument. Despite the fact that Oromos share long geographic boundaries with the Ethiopian Somali, the results in this study don’t show genetic relationship of the two better than that each has with others in the major Group 1. This might be due to the isolation of the sampling areas of the two communities. We still believe a strong genetic contact of the two along their boundaries. However, the contact between the two people is likely during the post 16^th^-century Oromo migration (Lewis 1966). Hence, their genetic relation may quickly decay as we go away from their current shared areas.

The other impressive result in this study is the genetic relationship of the Cushitic language speakers, Afar more with Amhara-Tigray than the other Cushitic language speakers: Oromo and Somali. The Afar and the Abyssinian highlanders have been in close contact since the Axumite time(Yasin 2008). The Abyssinian Christian rulers and Adal (Afar) Sultanates have been in long standing rivals and sometimes allies in the horn of Africa. However, Afar has more intimate cultural, religious, and physical relationships with Somali than the Abyssinian highlanders, which violates the clear genetic relationship more with Abyssinian highlanders than both the Ethiopian Somali and the Somali in mainland Somalia. Perhaps the Afar people are close relatives and/or descendants of the Amhara Shewa (Trimingham 1952). These two were with overlapping rulers for centuries between the Shewa Christian and Walasma (Ifat) Islamic dynasties. The Shewa Christian rulers went far to the portal center of the Ifat Sultanates Zeila; while the Ifat Sultanates also advanced well deep in the empire of the Christian kingdom up to the current Menz area in Shewa (Pankhurst 1997).

The haplotypes distribution possibly portrays the gene flows that are attributed to the historical movement of the people. HAP2 and HAP3, which tend to diminish as we go further south, were likely spread following the first North-South movement. This also may entail the initial relationship of all the people in the horn of Africa, which mostly was under the influence of the Abyssinian and Walasma empires in the region (Pankhurst 1997). HAP10, which has spread among the populations within the Ethiopia boundary, might be due to recent admixture events. This haplotype likely originated from the south Omotic, the Aries that perhaps started to spread since the Northerners met the Aries. Interestingly, this haplotype is absent in Somali from the Somali mainland, which may further strengthen our argument of its recent success into the north. The HAP10 is also seen faintly among Nilotics that the highland Christian empires rarely expanded into. We see its gradient as we move from Gumuz, which has a long history under the highland Christian rulers, Anuak, that recently joined Ethiopian and the South Sudanese who share borders with the Anuak, which is still against the spatial distance of these populations to the Ari, the likely origin of the HAP10.

The genetic differentiation among the Ethiopian populations is higher than populations living in a wider geographic region elsewhere(Biswas et al. 2009; Nelis et al. 2009). This might be due to un-melted ancestral lines of different origins living in the region, forming a multi-ethnic mosaic society (Levine 1974). The genetic complexity of Ethiopians, which attributes to historical gene-flow to the region, has been demonstrated in several recent studies (Hassan et al. 2008; Lovell et al. 2005; Pagani et al. 2012; Passarino et al. 1998). Ethiopia and its surroundings probably harbor a higher genetic diversity than most parts of the world (Pagani et al. 2012). The expected heterozygosity of Ethiopians is relatively high and matches the one reported in Hollfelder et al. (2017).

### The genetic landscape of Ethiopians against others

The high within Ethiopian genetic variation (Table 1) and hence high genetic diversity (Fig 5) compared to others is in line with several previous reports on Ethiopian genetic lineage. Earlier studies showed ~a 40-50% non-African genetic composition of most Ethiopian and the African horn populations (Lawson et al. 2018; Pagani et al. 2012). In this study, with the subset dataset of Pagani et al. (2012) and CEU as an non-Africa reference ancestry, Ethiopians: Afar, Amhara, Tigray, and most Oromo contain 10-15% CEU (Fig 4). The disparity with the previous study might be due to the difference in reference out of Africa ancestral population, which may mask part of the admixture within African horn (see brown in Fig 4B and dark gray in Fig 5). However, in line with the previous studies (Gurdasani et al. 2015; Pagani et al. 2012; Pagani et al. 2015), Ethiopians show high genetic diversity (Fig 6), harboring the genetic breadth of all the target populations (Fig 4B; Fig 5) which is attributed to the immigration of the people from different geographic regions of the world (Llorente et al. 2015; Phillipson 1993).

The close relationship of the MKK (Maasai) with the most Ethiopians might also be attributed to the historical expansions and migrations of people. The Maasai claim their origin from the lower Nile and migrated south through Lake Turkana (Kihara 2020), the largest the Great Rift Valley lake that stretches from Northern Kenya to Southern Ethiopia (Kolding 1992). Maasai probably had contacts with Ethiopians through their movements in the regions. The results from Pagani et al. (2012) previously showed a predominant ancestral line of Maasai that spread all throughout Ethiopians except in Ari Blacksmith. This ancestral line was also seen in Egyptians, which might bear out the oral history of the Maasai that claims their origin from the lower Nile (Kihara 2020). Interestingly, the Maasai, given their historical geographic proximity to Ari blacksmith via Turkana, they are genetically less related despite the fact that Ari blacksmith ancestral line has spread in all Ethiopians, including the northern highlanders. The noticeable European ancestral line in this study and even higher ‘non-African’ admixture of Pagani et al. (2012) may invite a further investigation on historical contacts of the Maasai with others in the region. The comparable high gene diversity in Maasai and the Ethiopians in major group 1, unlike others, including the indigenous Africans, probably entails the historical relationships of the Maasai with the Ethiopian northerners. The Maasai might be a remote decedent of the Ethiopian northerners. Ari blacksmith probably met the Ethiopian in major group 1 after the Maasai left Ethiopians. Maasai might then met other ancestral lines such as Yoruba, where they went into (Pagani et al. 2012). The low population gene diversity in Ari Black Smith perhaps indicates inbreeding which probably is associated with the isolation of this community due to their occupation (Pagani et al. 2012; van Dorp et al. 2015).

### Allelic distribution of potentially adaptive loci and implication to success of people into geographic regions

Allelic variation among the populations at some loci hints natural selections that have relevance for local adaptation. This may hence determine the type of people who may have success in certain eco-geographic regions. For example, one of the SNPs that have shown strong adaptive signals (Fig. 7, S4) rs10900588 on Chromosome 1 has recently been reported for its association with Malaria-driven selection in *ATP2B4* gene (Gouveia et al. 2019). The higher frequency of G in the Nilotics: South Sudan, Anuak and Gumuz is also consistent with their finding that has demonstrated the high frequency of the G allele in Nilotic predominant North Uganda than the southern Uganda where dominated by Bantus. Our finding is also precisely in line with the resistance of the Nilotic to malaria disease compared to the rest of Ethiopians regardless of age and personal conditions (Armstrong 1978; Mathews and Armstrong 1981). The G allele, which is predominant in Nilotics at this locus is nearly absent in other Malaria infested inhabitants of the Northeast lowlands such as Afar and Somali that are geographically isolated from the Nilotics while this allele is at a traceable frequency in populations that have contact with the Nilotics (Fig 7, S4). The high frequency of this allele in YRI may also suggest the Malaria resistance in the Yoruba population (Network et al. 2014). The other outlier loci that showed a strong signal of adaptation can also be associated with certain phenotypic features. We suggest further studies of these loci, particularly the consistently high peak SNPs chromosome 23 and 24 (Fig S1-3).

## Acknowledgements

We are grateful to Luca Pagani for his help to access Ethiopian genotype data.

## Data availability

Genotype data of the Ethiopian dataset can freely be downloaded from http://mega.bioanth.cam.ac.uk/data/Ethiopia/

Genotyped data of CEU, MKK and YRI are available at ftp://ftp.ncbi.nlm.nih.gov/hapmap/genotypes/hapmap3_r3/

Linguistic tree data can be downloaded from https://osf.io/cufv7/files/

