## Supplementary for "The genetic and linguistic structures of Abyssinians and their neighbor reveal the historical demographic dynamics and environmental adaptation in the African Horn region" (from another version of this preprint: version 1, 2020-10-22)

**Supplementary Table**

Table S1 Top 20 outlier loci from the merged dataset, linked gene and associated health risks

| SNP_ID | Chr | Pos | Pval5 | Pval6 | Pval7 | Gene | Disease |
|---|---|---|---|---|---|---|---|
| rs7746960 | 6 | 52470824 | 3.44e-96 | 1.31e-94 | 3.19e-95 | TRAM2 | |
| rs1217384 | 1 | 114221283 | 4.31e-96 | 2.81e-95 | 1.42e-95 | AP4B1-AS1, BCL2L15, PTPN22 | Diabetes |
| rs17013967 | 4 | 89390337 | 3.70e-92 | 1.00e-91 | 1.77e-96 | | |
| rs3749820 | 5 | 132560962 | 1.70e-91 | 2.261e-90 | 8.91e-93 | FSTL4 | Iluem Cancer |
| rs1045737 | 9 | 134559562 | 4.9e-90 | 3.82e-89 | 3.760e-92 | GTF3C4 | |
| rs10804967 | 4 | 5769597 | 6.32e-90 | 1.19e-88 | 2.04e-91 | EVC | |
| rs2353193 | 2 | 170142309 | 6.7e-89 | 2.09e-88 | 5.00e-90 | | |
| rs739869 | 20 | 51303743 | 1.126e-86 | 1.8 4e-85 | 8.079e-89 | TSHZ2 | |
| rs16864613 | 3 | 154038142 | 2.51e-85 | 5.06e-84 | 2.73e-87 | PSD3 | |
| rs3758143 | 8 | 18434303 | 6.38e-84 | 4.63e-83 | 5.61e-88 | | |
| rs12947218 | 17 | 39987396 | 2.69e-79 | 8.39e-78 | 1.00e-76 | | |
| rs194071 | 5 | 9737199 | 4.81e-78 | 2.50e-77 | 2.04e-80 | LINC02112 | |
| rs2073716 | 6 | 31230976 | 5.40e-78 | 4.19e-77 | 1.22e-80 | CCHCR1 | |
| rs10782838 | 1 | 92596186 | 8.81e-75 | 2.00e-73 | 1.6 e-75 | RPAP2 , GLMN | |
| rs10793124 | 11 | 75129915 | 1.51e-72 | 3.21e-71 | 3.15e-70 | LOC105369393 | |
| rs12033610 | 1 | 225816475 | 8.90e-72 | 2.28e-70 | 8.62e-71 | ZNF678 | |
| rs6185 | 8 | 25336717 | 1.28e-69 | 4.87e-69 | 2.00e-73 | GNRH1 | Breast Cancer |
| rs10222558 | 3 | 31581784 | 4.88e-68 | 1.0e-66 | 1.02e-66 | STT3B | |
| rs9784858 | 6 | 32895153 | 4.50e-67 | 7.32e-66 | 6.30e-68 | | |
| rs11183722 | 12 | 45755490 | 1.64e-65 | 2.84e-64 | 7.47e-66 | AMIGO2 | |

Chr=Chromosome; Pos=SNP physical position on the chromosome, Pval5, Pval6, and Pval7= *P-value* at K=5, 6 and 7 respectively.

**Supplementary figures**

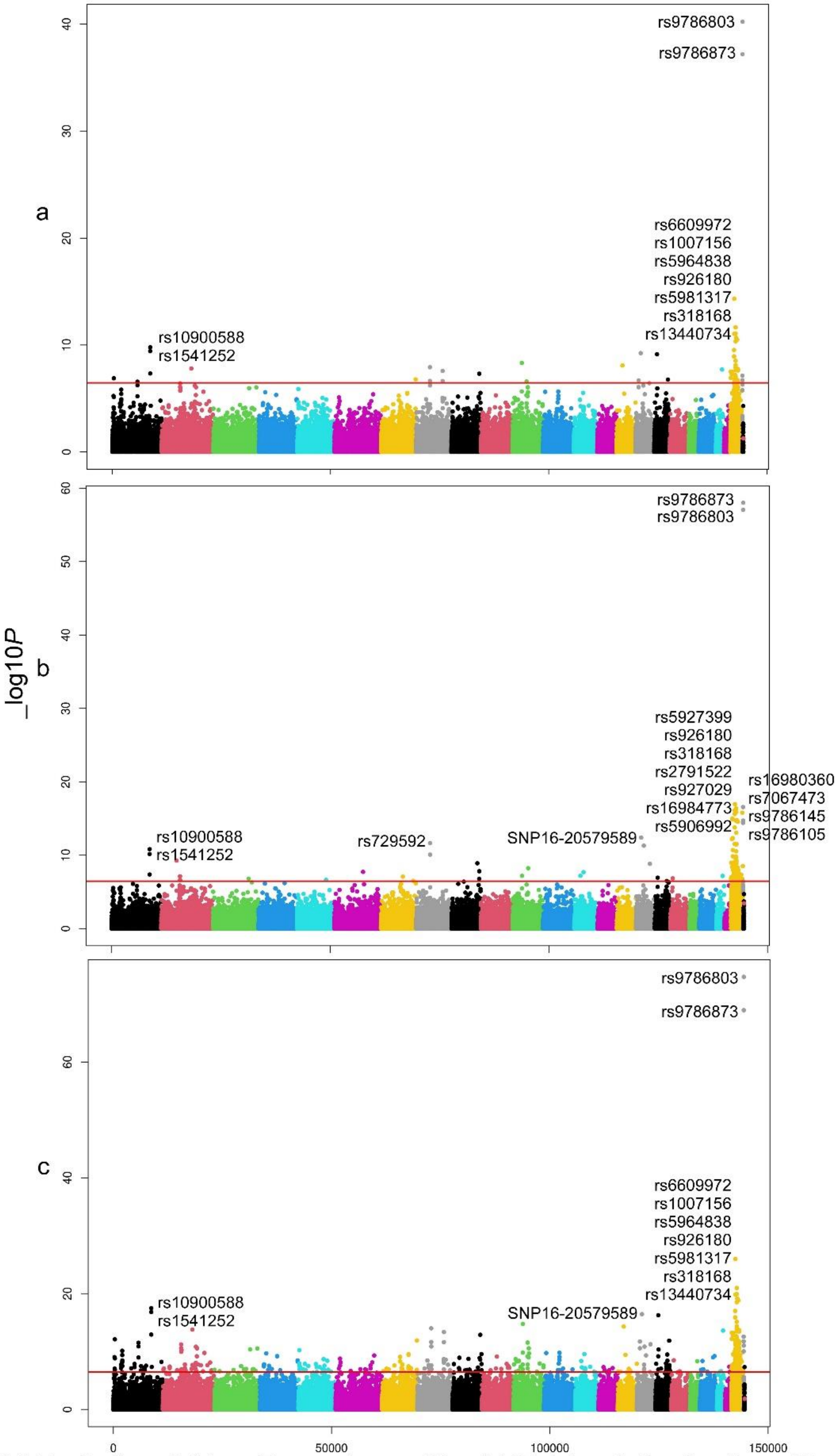


Fig S 1 Manhattan plot from $F_{ST}$ analyses of the 144K dataset showing the outlier loci a-c K 5, 8 and 12 respectively

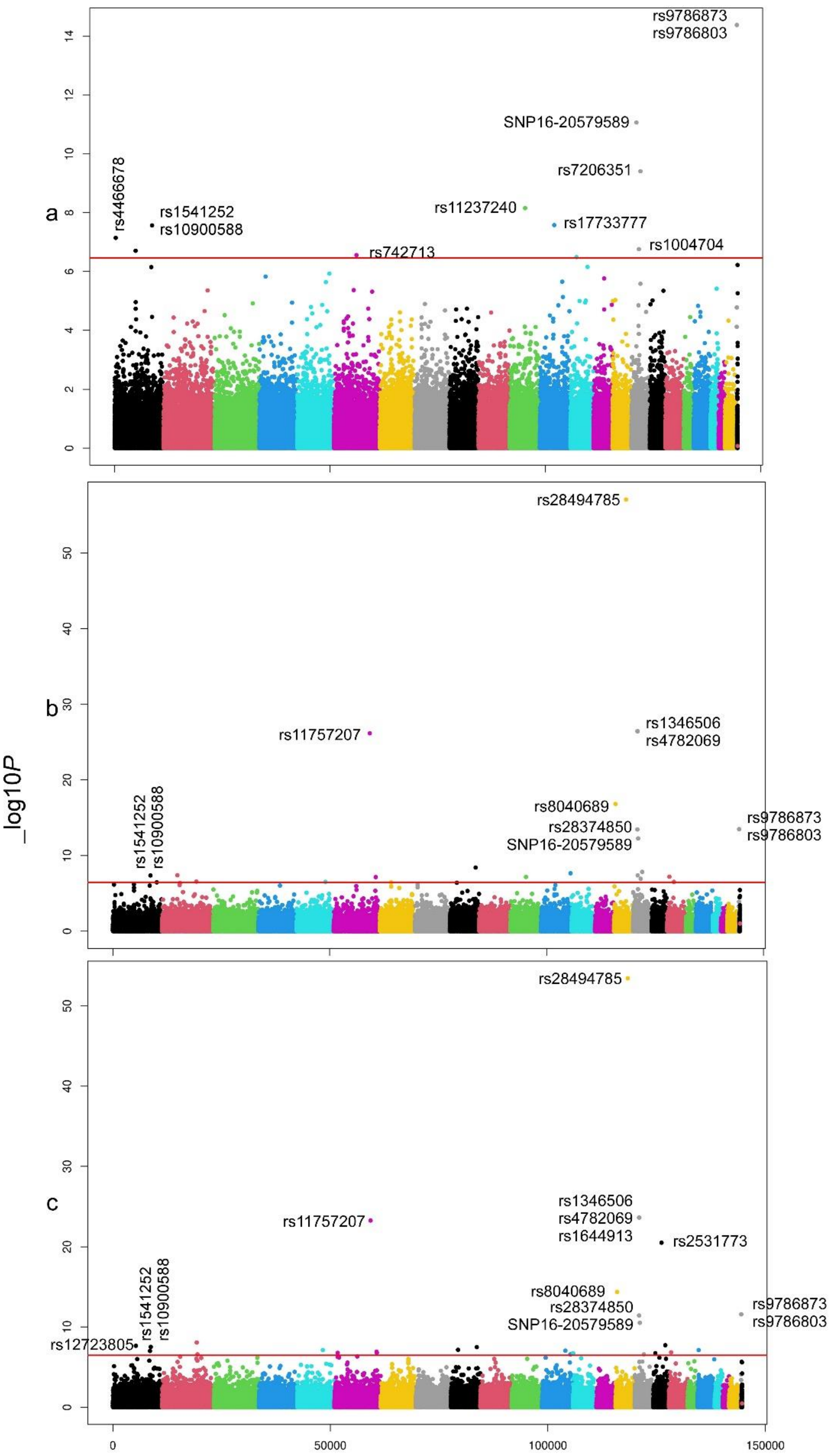


Fig S 2 Manhattan plot from pcadapt analyses of the 144K dataset showing the outlier loci a-c K 5, 8 and 12 respectively

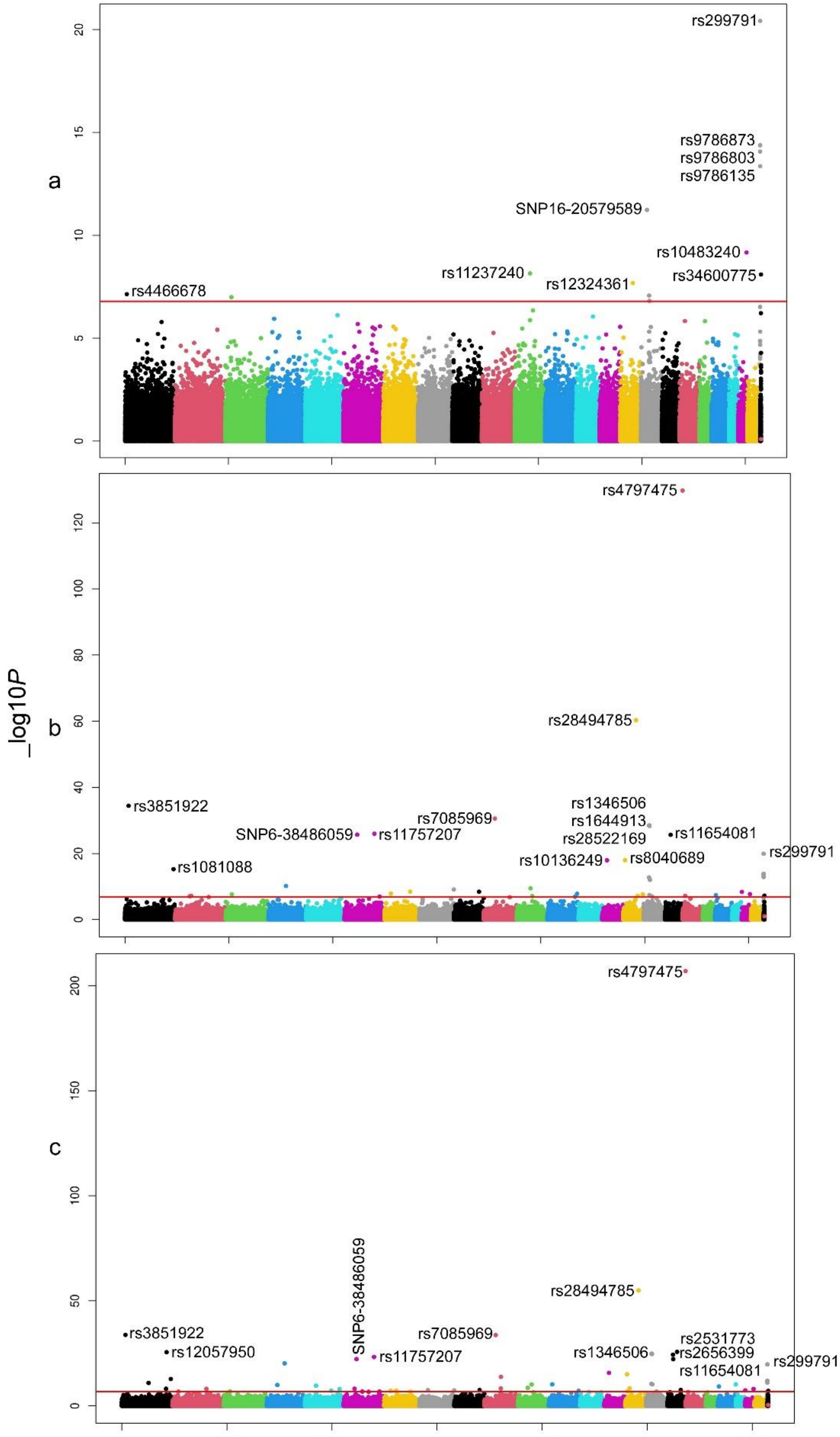


Fig S 3 Manhattan plot from pcadapt analyses of the 300K dataset showing the outlier loci a-c K 5, 8 and 12 respectively

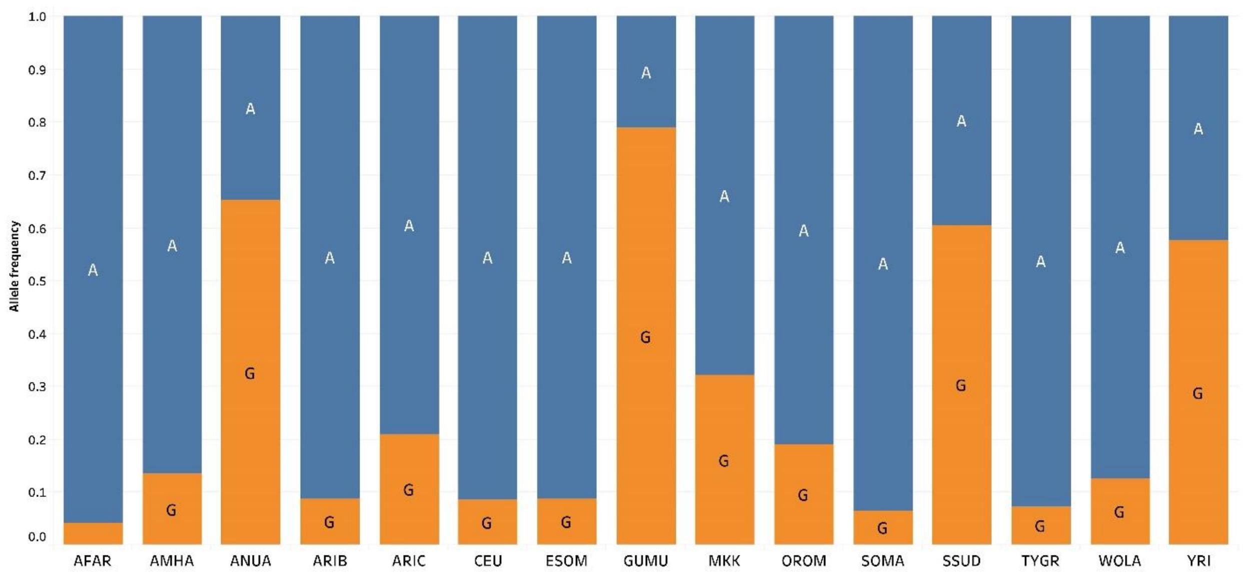


Fig S4 Allele frequency of Rs10900588 among populations
